# Using synthetic datasets to better understand and explain health outcomes associated with common single nucleotide polymorphisms

**DOI:** 10.1101/765586

**Authors:** Thomas R. Wood, Nathan Owens

## Abstract

Due to decreasing costs and a move towards “personalised medicine”, the use of direct-to-consumer genetic analyses is increasing. Both consumers and healthcare practitioners must therefore be able to understand the true disease risks associated with common genetic single nucleotide polymorphisms (SNPs). However, most population studies of common SNPs only provide average (+/−error) phenotypic or risk descriptions for a given genotype, which hides the true heterogeneity of the population and reduces the ability of an individual to determine how they themselves might truly be effected. Here, we describe the use of synthetic datasets generated from descriptive phenotypic data published on common SNPs associated with obesity, elevated fasting blood glucose, and methylation status. Using both simple statistical theory and full graphical representation of the generated data, we show that single common SNPs are associated with a less than 10% likelihood of effecting final phenotype, even in homozygotes. The significant heterogeneity in the data, as well as the baseline disease risk of Western populations suggests that most disease risk is dominated by the effect of the modern environment.

## INTRODUCTION

Due to decreasing costs and a move towards “personalised medicine”, the use of direct-to-consumer (DTC) genetic analyses and third party interpretation services is increasing.^1^ Though whole genome sequencing (WGS) is also increasing in popularity, most DTC products involve the analysis of common single nucleotide polymorphism (SNPs). These SNPs are then reported, either by the testing company or a third party tool that analyses the data, with specific disease risks based on published population data such as that from genome-wide association studies (GWAS). These risk predictions are generally based on population average outcomes, with the heterogeneity of a given phenotype or disease risk infrequently reported. In fact, most GWAS studies tend to only report descriptive data (e.g. mean and standard error) for a given phenotype (such as body mass index, BMI, or fasting blood glucose) within a risk genotype. By only comparing or providing group averages based on genotype, the consumer is likely to overestimate the disease risk associated with a given SNP. Presenting only simplified descriptive data, either graphically or numerically, for a given genotype gives the impression that each SNP has consistent penetrance with respect to the phenotype in question, which is known to not be the case.^2^ Therefore, the interpretation of disease risk based on SNPs by those not involved in the original studies and without access to the original data is almost impossible.

More important than the mean population effects of a given SNP or combination of SNPs that influence a common phenotype is the likelihood of a physiologically-relevant effect in a given individual. This includes the likelihood that there is no overall effect of genotype, particularly compared to common environmental factors that drive chronic disease risk in high income countries such as diet, sleep, and exercise. In order to allow for healthcare practitioners or self-interested parties to better understand the likelihood of a given phenotype being altered by a specific genotype, we developed a method by which synthetic datasets could be generated and analysed. This is largely possible due to the fact that the effects of SNPs on measurable phenotypes are generally considered to follow a normal distribution, with the number of alleles or weighted genetic scores being linearly associated with the target phenotype. Using this approach, the significant heterogeneity of population data can be better understood, particularly with respect to how a given individual may or may not display phenotypic changes based on the presence of common genotypes.

## METHODS

### Selection of representative SNPs

One of the authors (TRW) performed at-home SNP analysis using a 23andMe DTC kit (23andMe, Mountain View, CA). The data were run through a third-party analysis tool (FoundMyFitness Genetic Report) to identify SNPs most commonly reported to be associated with differential disease risk. Individual studies and meta-analyses of per allele effects for common SNPs most strongly-associated with risk of type 2 diabetes (Melatonin Receptor 1B, MTNR1B rs10830963), obesity (Fat mass and obesity-associated protein, FTO rs9939609), and altered methylation and nutrient handling resulting in elevated homocysteine levels (Methylenetetrahydrofolate Reductase, MTHFR rs1801131 and rs1801133) were identified from the third party tool output as well as the online SNP wiki SNPedia.com. Due to the significant effect of ethnicity on SNP disease penetrance, example population data that were likely to most closely match the Anglo-Scandinavian background of the author were used, including data from deCODE (Iceland) and the Northern Finland Birth Cohort (NFBC) that were included in large multi-population GWAS studies.^3,4^ According to recently-published methods suggested by Pontzer *et al*., published hunter gatherer data for fasting glucose were used to provide an estimate of the effect of the Western environment on fasting glucose and diabetes risk compared to a published genetic risk score.^5^

### Generation of synthetic datasets

Published per allele or per genetic risk score means were used to construct synthetic datasets for a given phenotype. All publications assumed data were normally distributed and that per allele/genetic risk score effects were linear. If data were expressed as mean with standard error (SE) or 95% confidence interval (CI), the standard deviation (SD) was calculated using the number (N) of participants in each group, where SD=SE*√N and SD=√N*(width of 95% CI)/3.92. When the descriptive data were not included in the publication, as was the case for genetic risk scores associated with obesity and fasting blood glucose,^3,4^ they were estimated from published graphs by extracting images and determining the number of pixels in each column and error bar relative to the scale bars on the axes. In all cases, enough data was included in the manuscript body to confirm that at least one of the estimated values was correctly determined using this method (such as total number of participants, or mean values in the highest or lowest genetic risk groups). For each genotype and gene, 1,000 synthetic individuals were randomly generated to re-create a normally-distributed dataset with the same mean and SD characteristics as those in the associated publication. Numbers were generated using Python 3.7 and the NumPy (1.17.0) and Pandas (0.25.0) libraries. The necessary code is available on GitHub (https://github.com/root-causing-health/SNPGaussianDistGenerator). Visual inspection of the data (Prism version 8, GraphPad Software, San Diego, CA) confirmed that they were normally distributed.

### Statistical analysis

Each synthetic dataset was graphically represented using a violin plot to show the full distribution of the data. Percent chance of a null effect from a risk allele was calculated by determining the percent overlap of the normal distribution of the wild type phenotype with that of a risk genotype using statistics.NormalDist in Python 3.8 Beta. The percent likelihood of the phenotype in a risk allele group being at or below the mean value of the “wild type” was also calculated, and linear regression analysis was performed to determine the percent contribution of risk alleles to a given phenotype. Similar analyses were performed using published multi-SNP genetic risk scores for type 2 diabetes and obesity.^3,4^

### Alternative methods

To encourage attempts to perform similar analyses, a number of free online tools can be used that do not require significant technical skills. After calculating mean and SD as described above, free gaussian random number generators such as from Random.org (https://www.random.org/gaussian-distributions/) can be used to generate synthetic datasets. Though the Box-Muller transform used by this tool is unlikely to produce a truly normal distribution,^6^ this is also unlikely to meaningfully affect the outcome. Similar online tools can be used to determine the likelihood of being at, above, or below, a given point in a normal distribution to determine null effects of a given SNP or risk score (http://onlinestatbook.com/2/calculators/normal_dist.html). Finally, free online graphing software can be used to visually represent the datasets for visual examination of variability and overlap (https://plot.ly/), and perform linear regression analyses (https://www.graphpad.com/quickcalcs/linear1/).

## RESULTS AND DISCUSSION

### FTO rs9939609 (A:T) and risk of being overweight

Published meta-analyses suggest an increase in body mass index (BMI) of 0.3 kg/m_2_ per FTO rs9939609 A allele.^7^ From this meta-analysis, data from the NFBC at 31 years of age (n=4,435) were used as a graphical example (**Figure 1A**).^8^ Mean (SD) BMI across the three genotypes was 24.12 (3.87) kg/m_2_, 24.43 (3.94) kg/m_2_, and 24.82 (3.95) kg/m_2_ for TT, AT, and AA respectively. In this population the risk of being overweight (BMI >25 kg/m_2_) was 41%, 44%, and 48%, resulting in an absolute 7% increase in risk in the TT genotype. BMI at or below the TT genotype was 47% in those with the AT genotype, and 43% in those with the TT genotype. The likelihood of null effect (percent overlap in BMI distribution of those with AT and AA genotypes compared to TT) was 96.8% and 92.8%, respectively. Therefore, only 3.2% of AT and 7.2% of AA genotypes would be expected to display any increase in BMI due to FTO genotype relative to TT. Linear regression found a significant association between number of A copies and BMI (p=0.001, R_2_=0.0035), suggesting that only around 0.4% of the variability in BMI is determined by FTO genotype (**Figure 1B**).

**Figure 1A.**
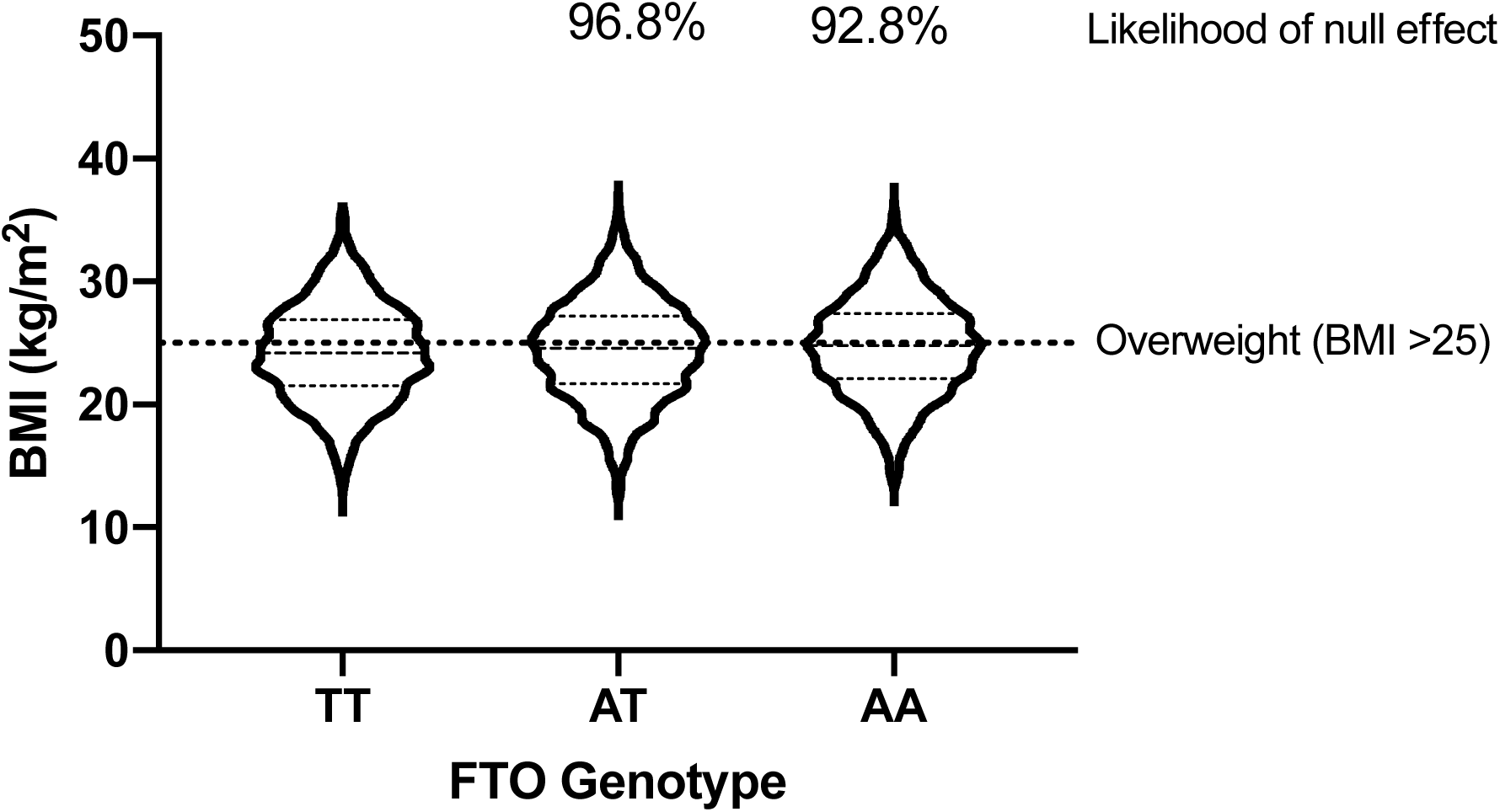
Effect of FTO rs9939609 genotype on BMI in the NFBC cohort. Violin plot displaying 1,000 synthetic BMI datapoints per FTO rs9939609 genotype, based on published population mean and SD values from the NFBC cohort.^8^ Percent overlap between the AT and AA normal distributions with that of the “wild type” (TT) genotype are displayed as a measure of the likelihood of these risk genotypes having no overall effect on BMI.

**Figure 1B.**
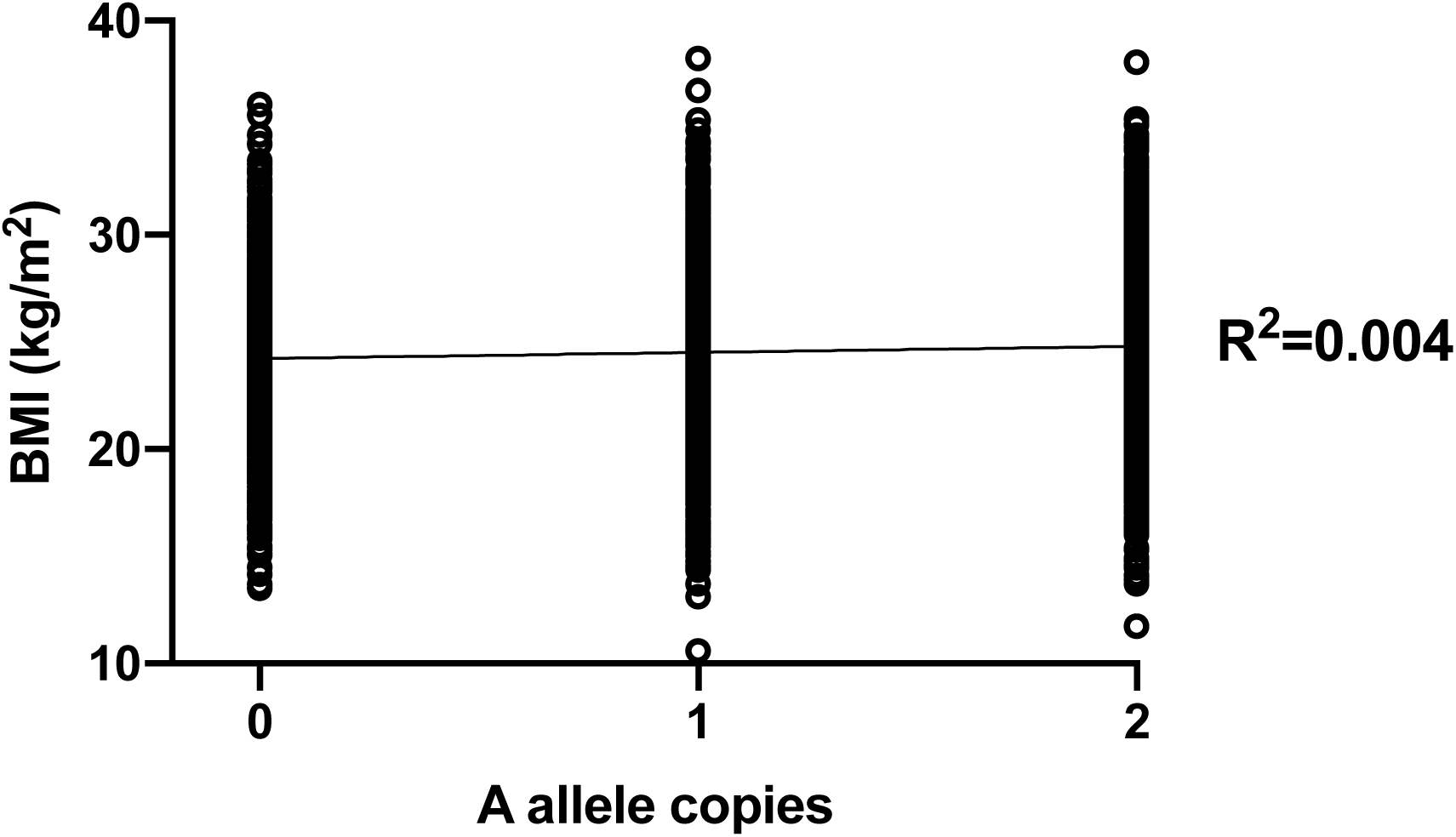
Linear regression of FTO rs9939609 A alleles versus BMI. Linear regression of 1,000 synthetic BMI datapoints per FTO rs9939609 A allele copy. There was a significant association between number of A copies and BMI (p=0.001, R_2_=0.0035), suggesting that only around 0.4% of the variability in BMI is determined by FTO genotype.

### Genetic BMI risk score

Willer *et al*. established a BMI genetic score using eight validated SNPs associated with BMI, weighted to effect size (with FTO rs9939609 given the largest weighting).^4^ This score was applied to the European Prospective Investigation of Cancer (EPIC) Norfolk cohort, where the top 1.2% of people (risk score >12) had an average BMI of 1.46 kg/m_2_ greater than those in the bottom 1.4% (risk score <4). However, the majority of participants had risk scores in the middle of the range (6-10), with large variability across the whole range of scores (**Figure 2A**). In the highest genetic risk groups (genetic scores of 11, 12, and >12), the likelihood of null effect was at least 80% (**Table 1**). The likelihood of null effect in the most common genetic score (score of 8, 18.4% of participants) was 88.1%. This suggests that regardless of an individual’s genetic score, there is less than a 20% chance that they will display any increase in BMI due to their score relative to those 1.4% of individuals with the lowest genetic risk. Across the entire range of scores, linear regression found a significant association between risk score and BMI (p<0.001, R_2_=0.018), suggesting that only around 2% of BMI is determined by the eight SNPs most significantly associated with BMI (**Figure 2B**).

**Table 1.**
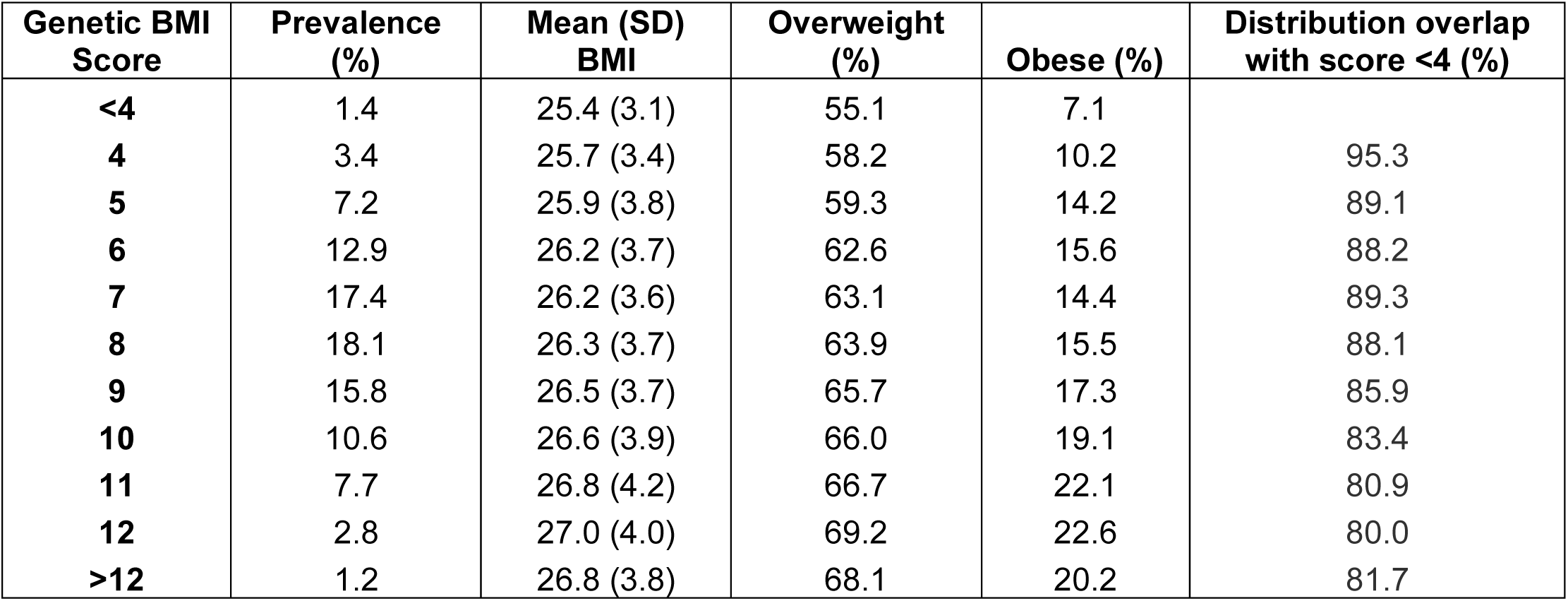
Effect of BMI genetic score on risk of overweight and obesity. BMI genetic risk score, as developed by Willer *et al*.,^4^ and risk of being overweight or obese, using population mean and SD values from the EPIC Norfolk cohort. Genetic scores of 6-10 cover around 75% of the population. The likelihood of null effect of each score was determined as the percent overlap of its normal distribution with that of the lowest risk group (score <4). Even in the highest risk groups (11, 12, <12) percent overlap was at least 80%, with only 12-17% of those with a genetic score of 6-10 predicted to have BMI affected by their genotype.

**Figure 2A.**
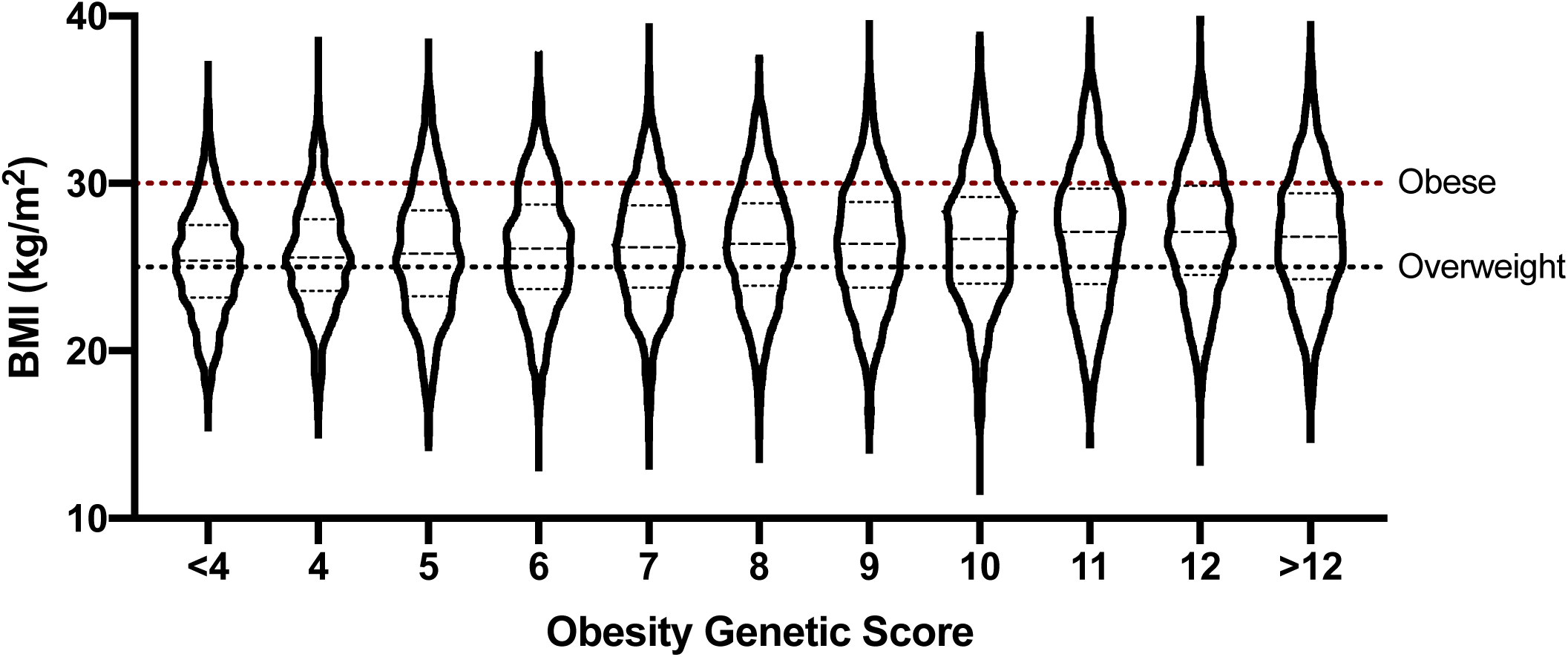
Effect of BMI genetic score on BMI in the EPIC Norfolk cohort. Violin plot displaying 1,000 synthetic BMI datapoints per group of BMI genetic risk score, as developed by Willer *et al*.,^4^ using population mean and SD values from the EPIC Norfolk cohort. Significant variability is seen across the entire range of genetic scores, with more than 50% of individuals being overweight regardless of genotype.

**Figure 2B.**
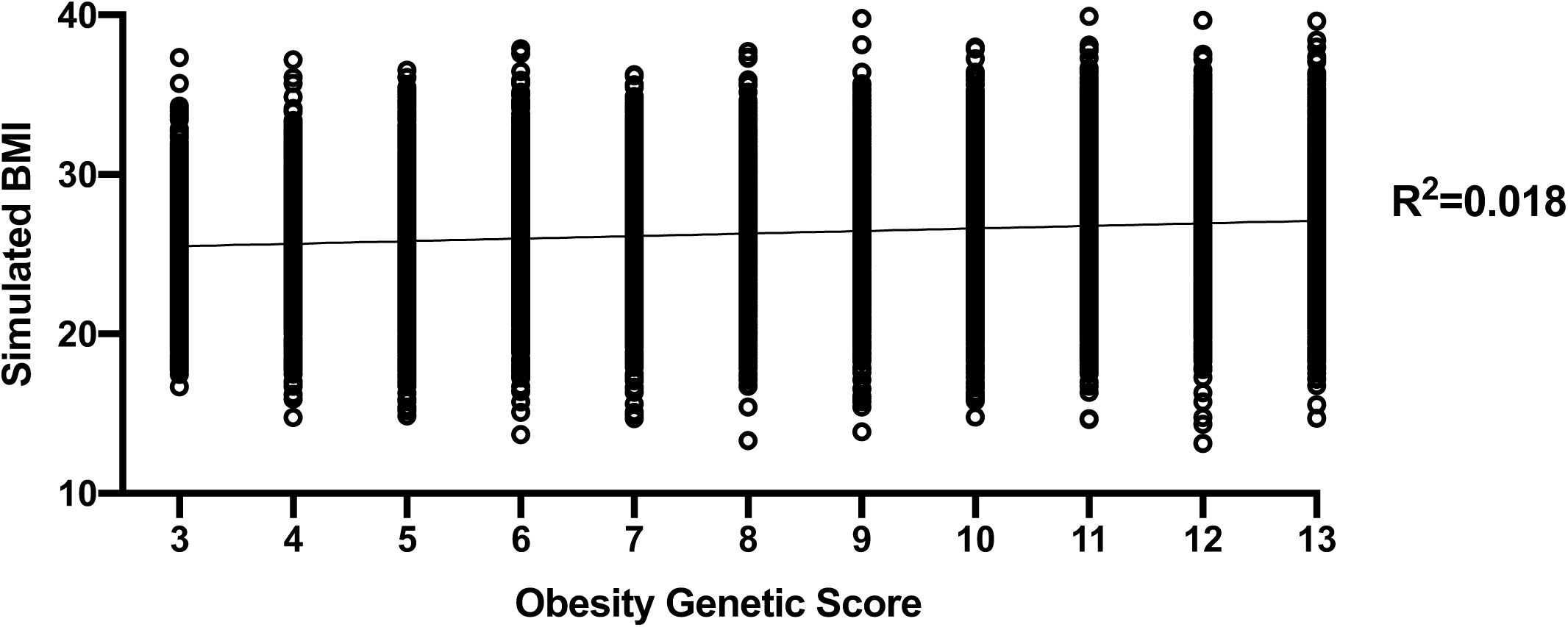
Linear regression of genetic BMI risk score versus BMI. Linear regression of 1,000 synthetic BMI datapoints per group of BMI genetic risk score, as developed by Willer *et al*..^4^ There was a significant association between risk score and BMI (p<0.001, R_2_=0.018), with around 2% of BMI determined by the eight SNPs most significantly associated with BMI.

### MTNR1B rs10830963 (C:G) and fasting blood glucose

Of the common SNPs associated with increased blood sugar, rs10830963 (C:G) has one of the largest effect sizes, with each G copy associated with around a 1.3 mg/dl increase in fasting blood glucose.^9^ Data from the deCODE cohort (n=6,240) were used as a graphical example (**Figure 3A**).^9^ Mean (SD) fasting blood glucose across the three genotypes was 95.2 (12.8) mg/dl, 97.0 (12.8) mg/dl, and 97.9 (12.8) mg/dl for CC, CG, and GG respectively. The likelihood of null effect was 94.4% in those with the CG genotype, and 91.6% in those with the GG genotype. Linear regression found a significant association between number of G copies and fasting blood glucose (p<0.001, R_2_=0.01), with around 1% of the variability in blood glucose being determined by MTNR1B rs10830963 genotype (**Figure 3B**).

**Figure 3A.**
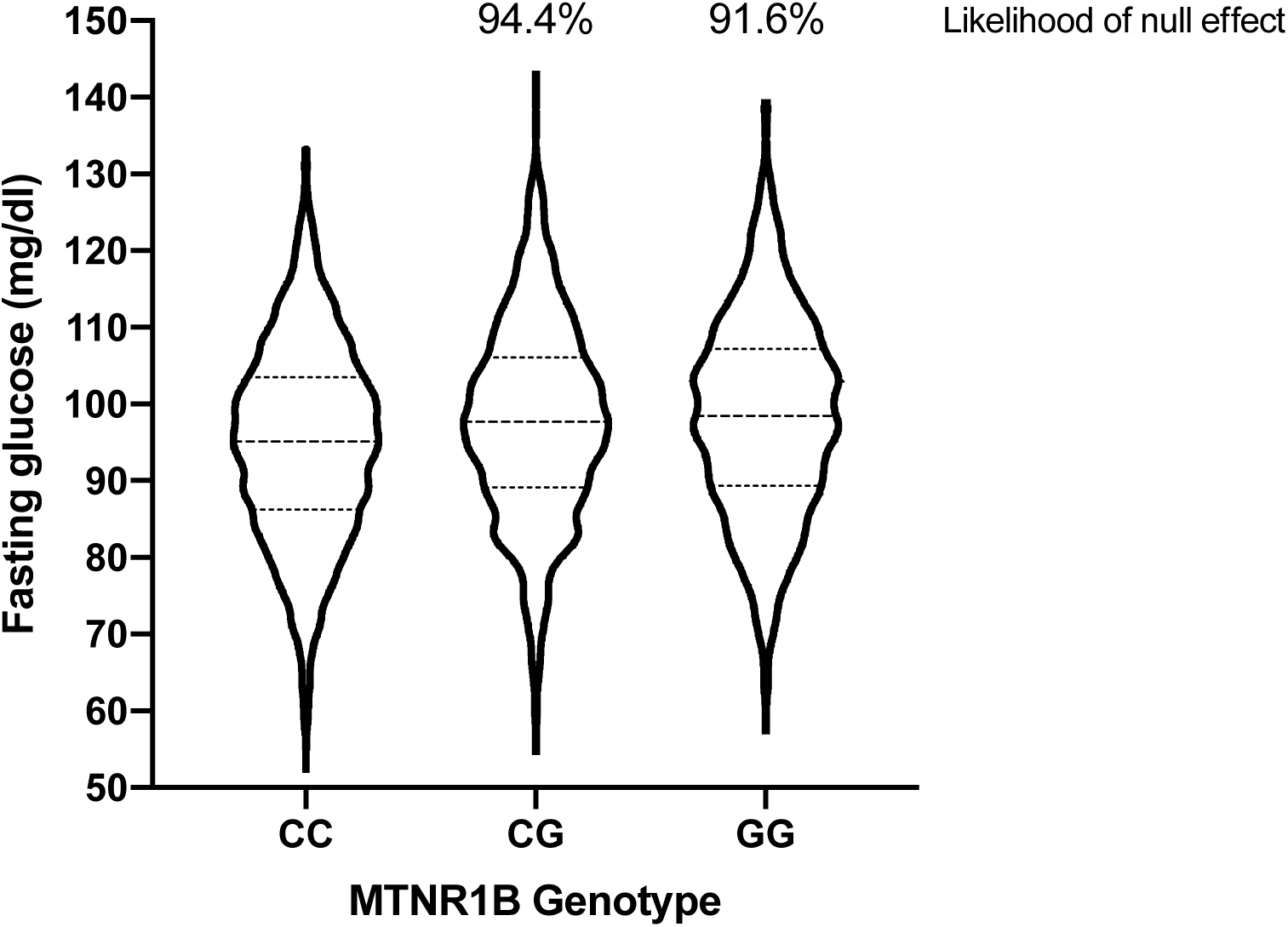
Effect of MTNR1B rs10830963 genotype on fasting glucose in the deCODE cohort. Violin plot displaying 1,000 synthetic glucose datapoints per MTNR1B rs9939609 genotype, based on published population mean and SD values from the deCODE cohort.^9^ Percent overlap between the CG and GG normal distributions with that of the “wild type” (CC) genotype are displayed as a measure of the likelihood of these risk genotypes having no overall effect on fasting blood glucose.

**Figure 3B.**
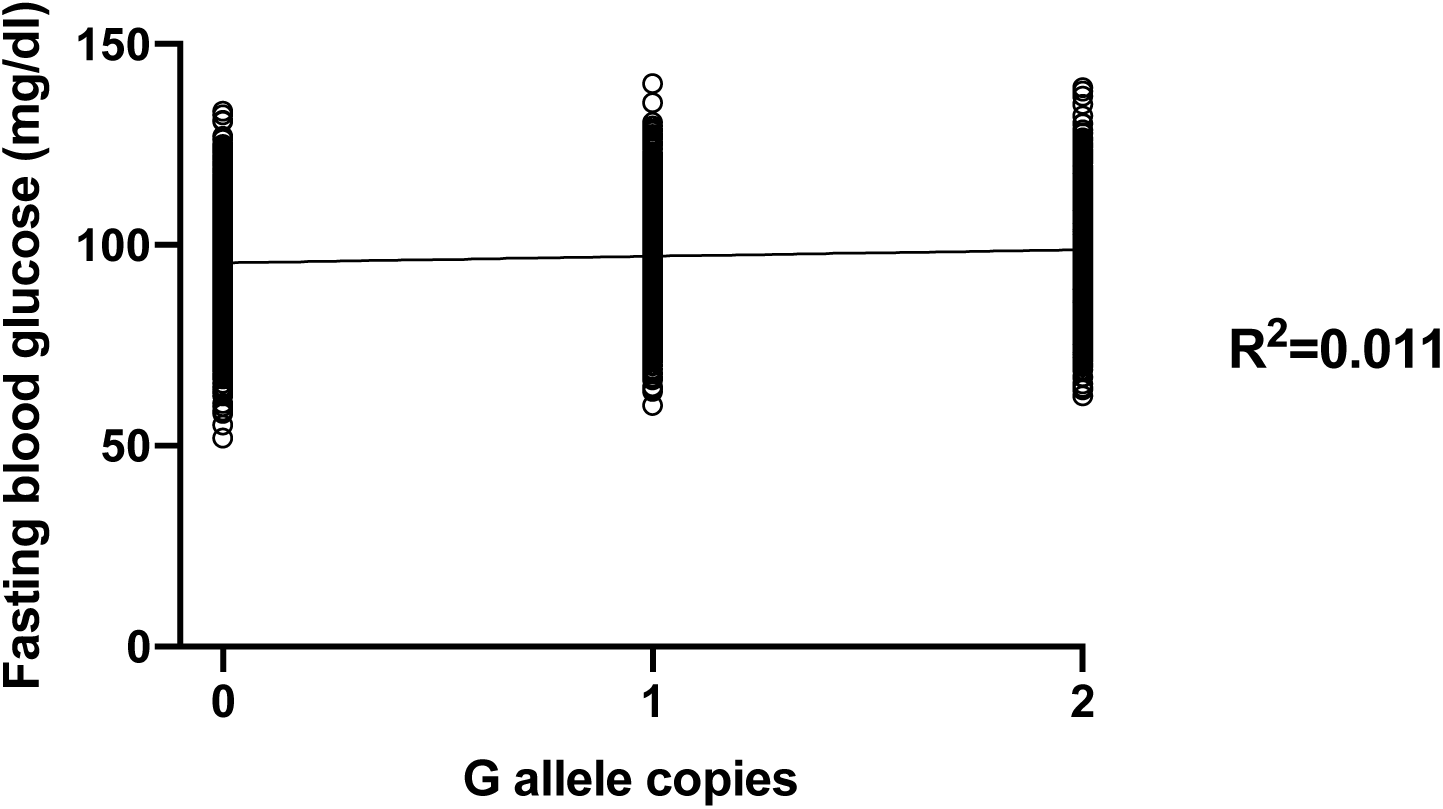
Linear regression of MTNR1B rs10830963 genotype versus fasting glucose. Linear regression of 1,000 synthetic BMI datapoints per MTNR1B rs9939609 G allele copy. There was a significant association between number of G copies and fasting glucose (p<0.001, R_2_=0.011), suggesting that only around 1% of the variability in fasting blood glucose is determined by MTNR1B genotype.

### Genetic type 2 diabetes risk score

Similar to the approach of Willer *et al*., Dupuis *et al*. published a genetic risk score for elevated fasting blood glucose and risk of type 2 diabetes,^3^ including MTNR1B and 15 other loci. This score was applied to the Framingham cohort, where the top 3.1% of people (risk score >22) had an average fasting blood glucose ∼6 mg/dl greater than those in the bottom 4.2% (risk score <13). Similar to the obesity risk score, significant heterogeneity in blood glucose levels was seen across the range of scores (**Figure 4A**). The likelihood of null effect in the most common genetic score (score of 18, 14.3% of participants) was 84.5% (**Table 2**). In those with the highest genetic risk score (scores 21, 22, and >22), the risk of prediabetic level blood glucose (>100mg/dl) was double that of those in the lowest risk group. However, even in these groups the likelihood of a given genetic score being associated with blood sugar outside of the distribution of those in the lowest risk group was only 25.5-27.7%, suggesting that fewer than 30% of people with the highest genetic risk of prediabetes experience that risk as a disease phenotype. Across the entire range of scores, linear regression found a significant association between risk score and fasting glucose (p<0.001, R_2_=0.049), suggesting that around 5% of fasting glucose is determined by the 16 SNPs most significantly associated with type 2 diabetes risk (**Figure 4B**). By comparison to the Framingham cohort, where mean (SD) fasting blood glucose was 92.5 (8.7) mg/dl in the lowest genetic risk group, free living hunter gathers from Tukisenta and Kitava reportedly have fasting blood glucose of around 75 (8) and 65 (14) mg/dl, respectively (**Figure 4C**).^10,11^ Based on these data, the Tukisentans would have a 98.6% likelihood of having a blood sugar below the mean of those in the Framingham cohort with the lowest genetic risk score, with a 97.5% likelihood in the Kitavans, and normal distributions that display only 19.5% and 27.3% overlap with the lowest risk Framingham group. This translates to a 0.09% and 0.05% risk of prediabetic fasting blood glucose, respectively. Therefore, even in the lowest risk genetic group in the Framingham cohort, the relative risk of prediabetic fasting blood sugar levels (19.4%) is around 200-400 times higher than in hunter gatherer populations.

**Table 2.** Effect of glucose genetic score on risk of prediabetes. Glucose genetic risk score, as developed by Dupuis *et al*.,^3^ and risk of having prediabetes, using population mean and SD values from the Framingham cohort. Genetic scores of 16-19 cover around 52% of the population, and have around 30% prevalence of prediabetes. The likelihood of null effect of each score was determined as the percent overlap of its normal distribution with that of the lowest risk group (score <13). In those with the highest genetic risk scores (21, 22, and >22), the risk of prediabetic blood glucose (>100mg/dl) levels was double that of the lowest risk group. However, even in these groups the likelihood of a given genetic score being associated with blood sugar outside of the distribution of those in the lowest risk group was only 25.5-27.7%, suggesting that fewer than 30% of people with the highest genetic risk of prediabetes experience that risk as a disease phenotype.

| Genetic Glucose Score | Prevalence (%) | Mean (SD) fasting glucose (mg/dl) | Prediabetes (%) | Distribution overlap with score <13 (%) |
| --- | --- | --- | --- | --- |
| <13 | 4.2 | 92.5 (8.7) | 19.4 |  |
| 13 | 5.0 | 93.6 (8.8) | 23.4 | 94.8 |
| 14 | 8.2 | 94.2 (8.6) | 25.0 | 92.3 |
| 15 | 9.8 | 94.3 (8.8) | 25.9 | 91.7 |
| 16 | 13.0 | 95.2 (8.9) | 29.5 | 87.7 |
| 17 | 13.8 | 95.4 (8.7) | 29.9 | 86.7 |
| 18 | 14.3 | 95.9 (8.9) | 32.3 | 84.5 |
| 19 | 11.5 | 96.5 (8.7) | 34.4 | 81.9 |
| 20 | 8.5 | 97.3 (8.8) | 38.0 | 78.3 |
| 21 | 5.4 | 98.1 (8.6) | 41.3 | 74.5 |
| 22 | 3.2 | 98.6 (8.7) | 43.6 | 72.7 |
| >22 | 3.1 | 98.6 (8.6) | 43.5 | 72.3 |

**Figure 4A.**
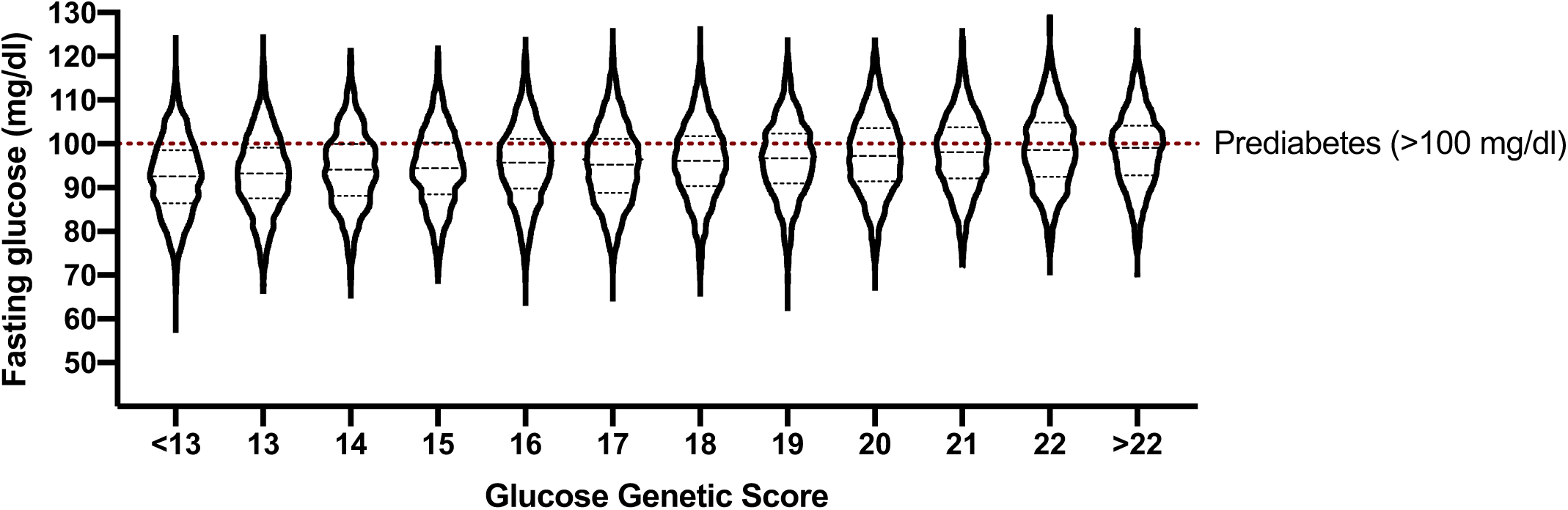
Effect of glucose genetic score on fasting glucose in the Framingham cohort. Violin plot displaying 1,000 synthetic glucose datapoints per group of glucose genetic risk score, as developed by Dupuis *et al*.,^3^ using population mean and SD values from the Framingham cohort. Significant variability is seen across the entire range of genetic scores. Risk of prediabetes (fasting glucose >100 mg/dl) increases from 19.4% to 43.5% from the lowest to highest risk group, with the most common genetic risk profiles (scores of 16-19, ∼50% of individuals) having around a 30% risk of prediabetes.

**Figure 4B.**
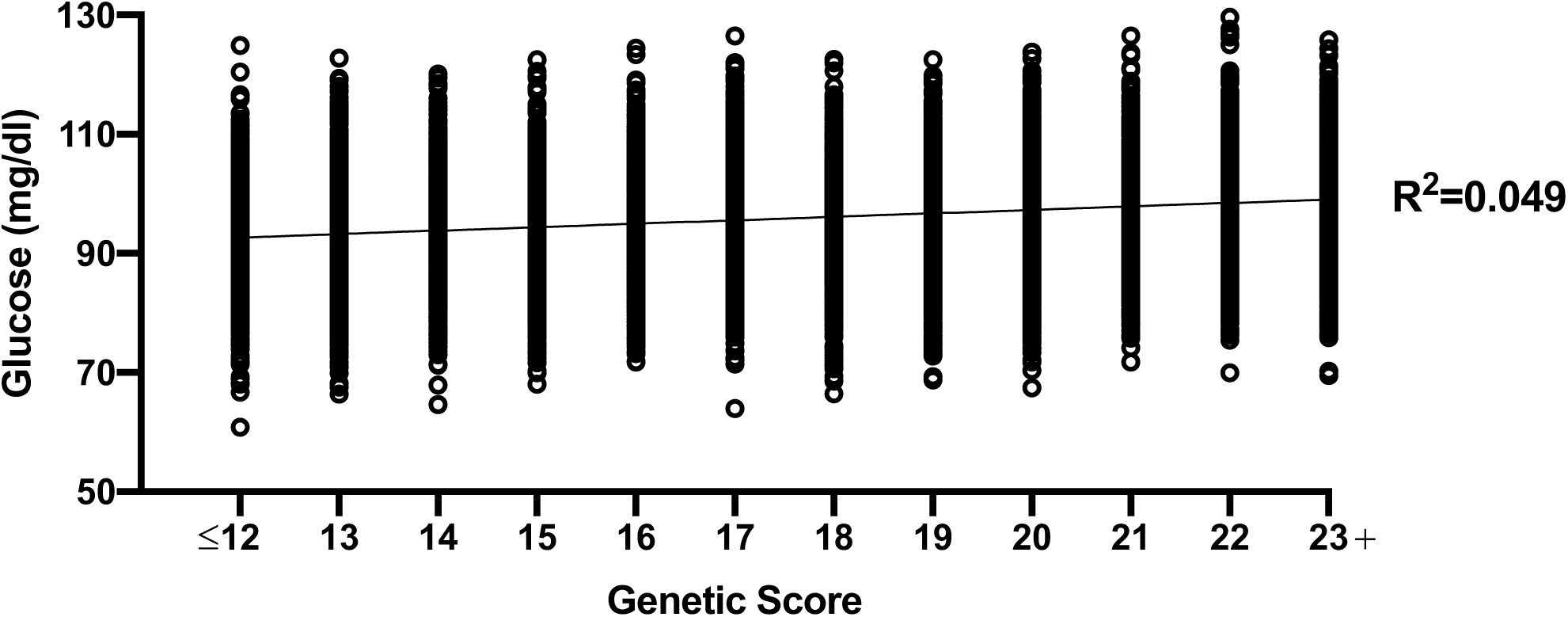
Linear regression of genetic glucose risk score versus fasting glucose. Linear regression of 1,000 synthetic BMI datapoints per group of glucose genetic risk score, as developed by Dupuis *et al*..^3^ There was a significant association between risk score and fasting blood glucose (p<0.001, R_2_=0.049), with around 5% of fasting glucose variability determined by the 16 SNPs most significantly associated with glucose homeostasis.

**Figure 4C.**
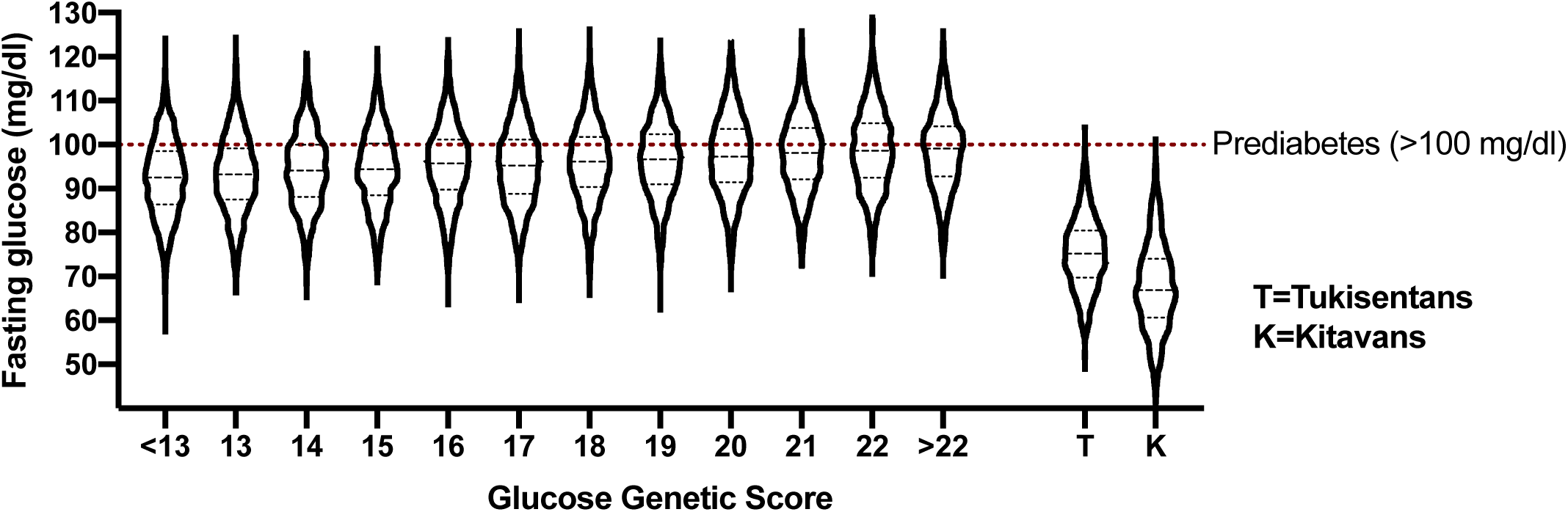
Comparison between fasting glucose in the Framingham cohort and in hunter gatherer populations. Violin plot displaying 1,000 synthetic glucose datapoints per group of glucose genetic risk score using population mean and SD values from the Framingham cohort,^3^ as well as using data from two hunter gatherer cohorts, the Tukisentans and Kitavans.^10,11^ The Tukisentans would have a 98.6% likelihood of having a blood sugar below the mean of those in the Framingham cohort with the best genetic score, with a 97.5% likelihood in the Kitavans. They display normal distributions with only 19.5% and 27.3% overlap with the lowest risk Framingham group and 0.09% and 0.05% risk of prediabetic fasting blood glucose, respectively. Even in the lowest risk genetic group in the Framingham cohort, the estimated prevalence of prediabetic fasting blood sugar levels (19.4%) is around 200-400 times higher than in hunter gatherer populations.

### MTHFR rs1801131 (A:C) and rs1801133 (C:T) and homocysteine

Two common polymorphisms in the MTHFR gene, which alter *in vitro* enzyme activity and are associated with reduced capacity to produce 5-methyltetrahydrofolate, are frequently discussed in the popular and alternative health fields with regard to the methyl cycle and associated changes in detoxification, cellular repair, and detoxification pathways. In 1998, van der Put *et al*. described *in vitro* MTHFR activity of the most common combinations of alleles at rs1801131 and rs1801133, as well as homocysteine levels in the same participants.^12^ In the most common genotypes, excluding 1298AA/677TT, which account for around 88% of the population on average, MTHFR function across five genotypes varies from 100% to 47.7% (**Table 3**). However, even in those with 47.7% function (1298AC/677CT) there is an 82.1% chance of null effect compared to 1298AA/677CC “wild type” with 100% function (**Table 3**). Across these common mutations, MTHFR function only explains around 1% of the variability in homocysteine levels (p<0.001, R_2_=0.01; **Figure 5A**). The addition of 1298AA/677TT, which has around 12% prevalence in the population and is associated with a 75.2% loss of MTHFR function, increases the explanation of variance to 7% (**Figure 5B**); however, the synthetic dataset included 6.9% negative values due to the large SD in this population. This suggests significant heterogeneity of homocysteine in those with the 677TT/1298AA genotype, which is not normally distributed. Indeed, though the percent chance of non-significant difference in homocysteine levels compared to 1298AA/677CC was only 35% in those with 1298AA/677TT, this includes a large proportion of the distribution in homocysteine levels that would be below that of the “wild type” due to the very large SD in the 1298AA/677TT group; 31.3% would be predicted to have homocysteine levels below the mean of 1298AA/677CC.

**Table 3.**
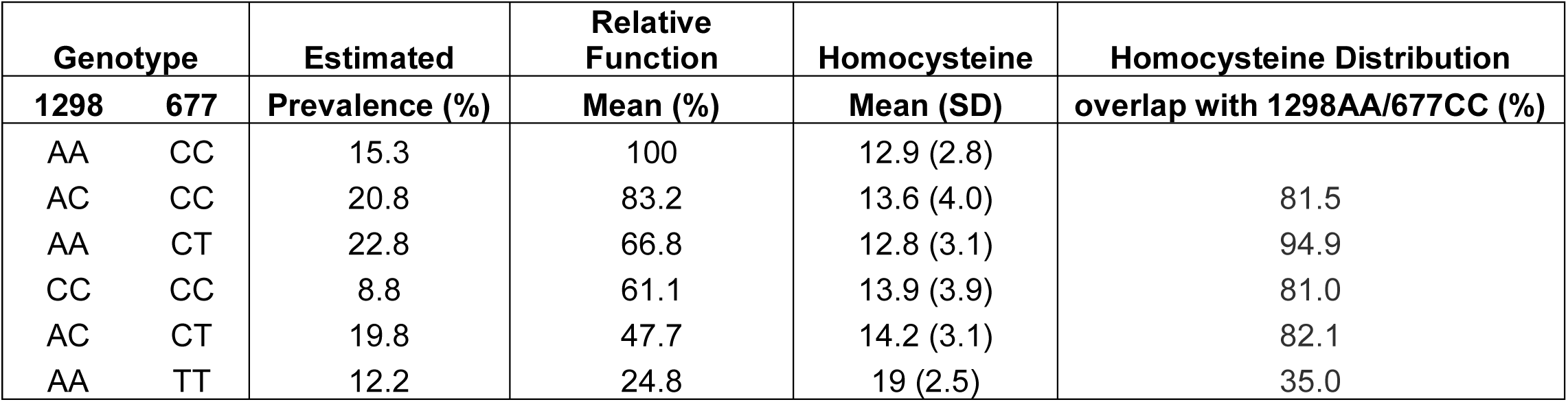
Effect of common MTFHR SNPs on average function and homocysteine. Average MTHFR function and homocysteine levels by rs1801131 (A1298C) and rs1801133 (C677T) SNP combination, as published by van der Put *et al*..^12^ Genotypes are listed in order of *in vitro* MTHFR enzyme function, with estimated population prevalence taken from Brown *et al*..^23^ The likelihood of null effect of each combination of MTHFR SNPs was determined as the percent overlap of its normal distribution with that with the “wild type” genotype (1298AA/677CC). Significant non-linearity between degree of MTHFR function and homocysteine is seen, with the most common genotypes, representing close to 88% of the population and 47.7%-83.2% enzyme function displaying 81-95% likelihood of null effect on resulting homocysteine levels.

**Figure 5A.**
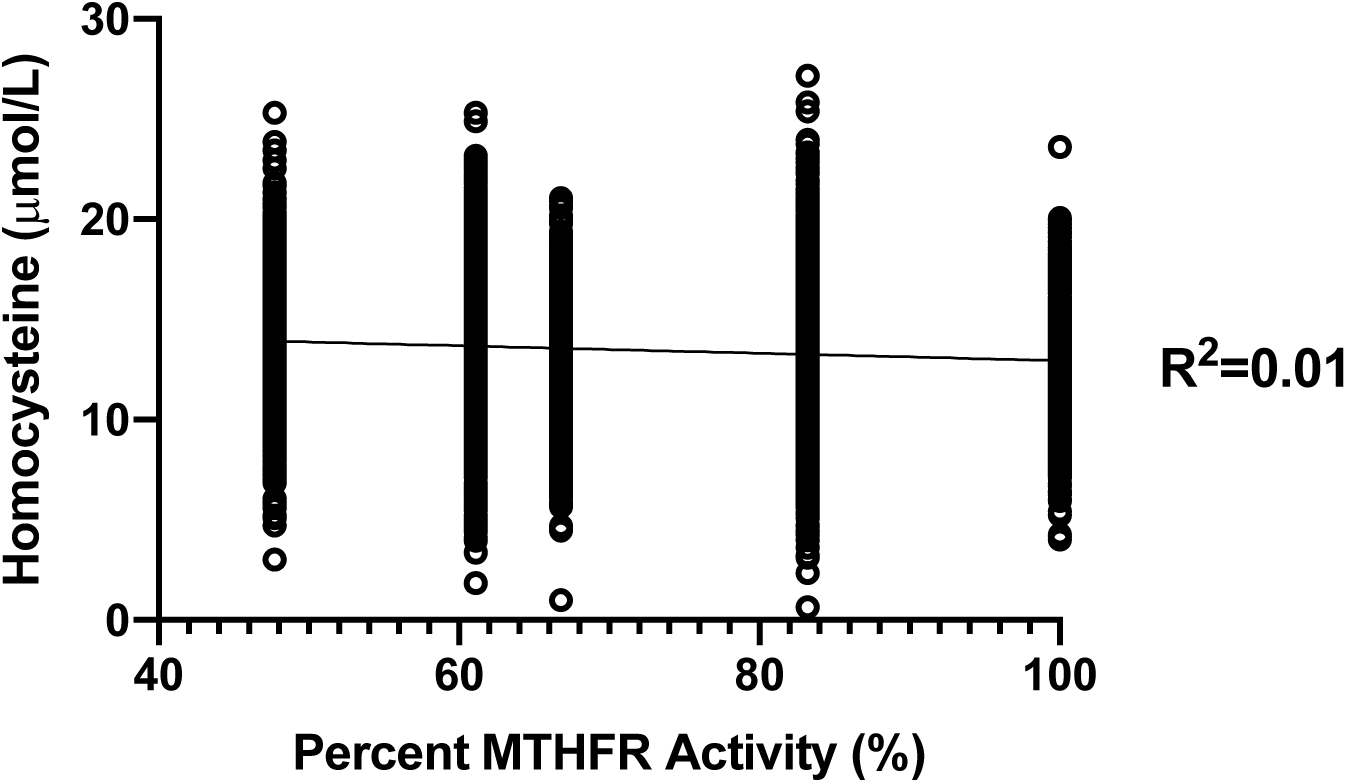
Linear regression of MTHFR activity versus homocysteine for the most common genotypes. Linear regression of 1,000 synthetic homocysteine datapoints per combination of common rs1801131 (A1298C) and rs1801133 (C677T) SNPs by *in vitro* MTHFR activity, excluding 1298AA/677TT. There was a significant association between MTHFR function and homocysteine (p<0.001, R_2_=0.01), suggesting that only around 1% of the variability in homocysteine is determined by MTHFR activity across these genotypes.

**Figure 5B.**
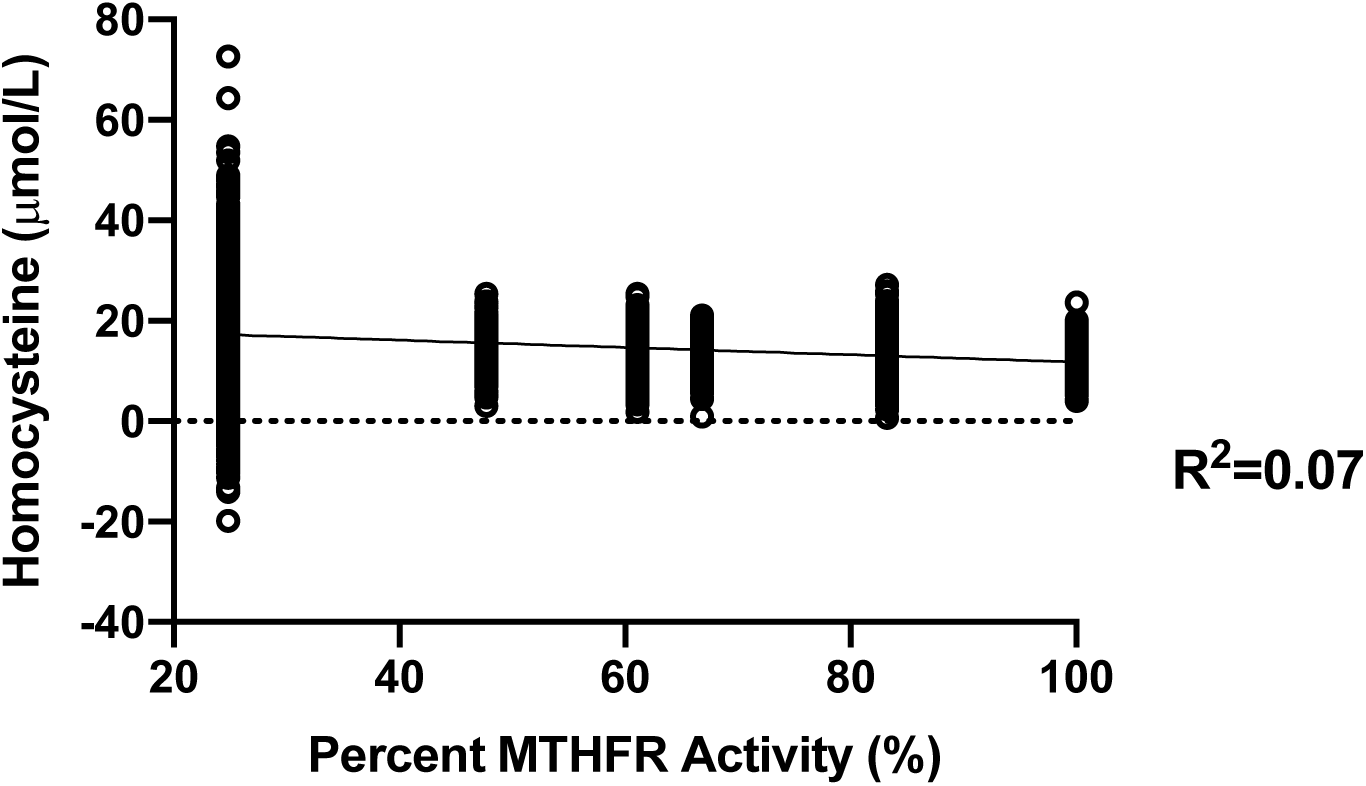
Linear regression of MTHFR activity versus homocysteine including 1298AA/677TT. Linear regression of 1,000 synthetic homocysteine datapoints per combination of rs1801131 (A1298C) and rs1801133 (C677T) SNPs by *in vitro* MTHFR activity. There was a significant association between MTHFR function and homocysteine (p<0.001, R_2_=0.07); however, the large SD (66% of the mean) in those with 1298AA/677TT resulted in 6.9% of predicted homocysteine levels being negative. This sugge sts that homocysteine in those with 1298AA/677TT is highly-variable, non-normally distributed, and that the effects of MTHFR activity on homocysteine levels are non-linear.

## DISCUSSION

The increasing prevalence of DTC genetic analyses is resulting in more and more healthcare providers being asked to interpret SNP-based disease risk by their patients, or attempting to incorporate these analyses into personalised treatment approaches. Here we demonstrate that, by using simple statistical theory and synthetic datasets generated based on published population phenotypic data from well-characterised SNPs, the likelihood of any given genotype resulting in a meaningful difference in phenotype is relatively small. For individual common SNPs determined to have large effect sizes, such as *FTO rs9939609* on BMI and *MTNR1B rs10830963* on fasting glucose, even those with two alleles have a less than 10% chance of displaying a difference in phenotype due to significant population variability. Additionally, baseline disease risks suggest that the vast majority of health outcomes associated with common SNPs are dominated by the environment.

The best-characterised SNP associated with risk of overweight and obesity is FTO rs9939609, with an average per A allele increase in BMI of 0.3 kg/m^2.7^ However, an average population effect is less useful to an individual than the likelihood that they are going to be affected in the first place. For a single FTO A allele, this likelihood is around 3%, increasing to 7% in individuals with two A alleles, with 0.4% of overall BMI explained by FTO genotype. Though it may be the SNP most well associated with increases in BMI, the vast majority of individuals are unlikely to have their BMI meaningfully affected by their FTO SNPs. Importantly, even this negligible effect of FTO on BMI is largely dominated by the environment, with recent analyses suggesting that FTO rs9939609 genotype was not associated with BMI in those born before 1942.^13^ Similarly, analyses of both FTO rs9939609 SNPs and composite obesity genetic risk scores suggest that those who partake in regular movement or exercise (∼1h of moderate-vigorous physical activity per day) have similar BMIs regardless of genetics.^14,15^ In the well-characterised EPIC Norfolk cohort, the risk of being overweight was above 50% regardless of a genetic score consisting of the eight SNPs most tightly-associated with BMI, again suggesting a significant environmental component. Considering that current Centers for Disease Control and Prevention data suggests that 39.8% of the adult population in the United States is obese,^16^ the degree to which common genetic SNPs contribute to BMI may be statistically significant but borderline physiologically irrelevant compared to the impact of the environment.

Similar results to those seen with genetic obesity risk were found when analysing genetic risk of elevated fasting blood glucose and type 2 diabetes. Of the SNPs associated with increased fasting blood glucose, MTNR1B SNP rs10830963 (C:G) has one of the largest effect sizes, with each G copy associated with around a 1.3 mg/dl increase in fasting blood glucose.^9^ In our analysis, only 5.6% of individuals with a single G copy would be expected to experience an increase in fasting blood sugar relative to those with the CC genotype, increasing to 8.2% in homozygotes. Using the genetic risk score developed by Dupuis *et al*. is more predictive, with more than a doubling of risk of prediabetes in those with the highest genetic frisk score compared to those with the lowest genetic risk. However, linear regression analysis suggested that only around 5% of fasting blood glucose is determined by genetic risk. This is just very similar to the proportion of explained variance that Dupuis *et al*. state in their original manuscript,^3^ which provides some support for the use of synthetic datasets when variance and absolute numbers are not provided in the published literature. More importantly, however, it the way in which this information is placed into the context of the consumer using DTC genetic analysis to assess disease risk. For instance, the variance in fasting blood glucose (∼5%) attributed to the loci included in the genetic risk score is smaller than the variance in reproducibility of commonly-used hand held at home glucometers used to monitor blood glucose in individuals with diabetes. Any effect of genetic risk is also largely a reflection of a slight amplification of the risk associated with the Western environment. Compared to hunter gatherer populations,^5,10,11^ fasting glucose is around 25-30 mg/dl higher even in the lowest genetic risk group, and the risk of prediabetes is 200-400 times higher. Indeed, in a recent analysis of the Bolivian Tsimane, prevalence of type 2 diabetes was 0%,^17^ on top of which any increase in genetic risk would be essentially meaningless. Therefore, the presence of any prediabetes appears to simply be a reflection of disease risk in the US as a whole, where more than 80% are thought to have suboptimal metabolic health, including more than 50% with fasting glucose >100 mg/dl.^18^ Based on multiple lines of evidence, close to 100% of the disease risk associated with elevated fasting blood glucose in the Western world can be attributed to the modern environment.

The concept of methylation capacity and its association with long-term health has recently gained a lot of interest in the alternative health community and popular press. As a result, DTC testing of common SNPs in the MTHFR and other related genes is being used to estimate an individual’s capacity to (re)generate methylfolate in order to guide disease risk or nutrient supplementation. One potential biomarker of methyl cycle function, including MTHFR activity, is homocysteine, which is associated with and increased risk of cardiovascular disease, dementia, and all-cause mortality when elevated.^19-21^ Though there are multiple pathways for the metabolism of homocysteine, one is dependent on methylfolate, and homocysteine levels are often used as a proxy for the status of the folate cycle.^22^ Importantly, SNPs resulting in decreased *in vitro* MTHFR function are common. The “wild type” genotype 677CC/1298AA associated with 100% MTHFR function is only found in around 15% of the population,^23^ which makes some degree of reduced MTHFR function a more representative “normal” state. In addition to this, the degree of MTHFR function appears to be only loosely associated with homocysteine levels. For instance, only 1% of homocysteine was accounted for by the five rs1801131 (A1298C) and rs1801133 (C677T) combinations that encompass 47.7-100% mean MTHFR activity. This suggests significant redundancy in the system that is unlikely to be able to inform any interventions based solely on genotype. Additionally, homocysteine levels are more likely to be determined by factors not associated with direct enzyme function, as those with the 1298AC/677CC genotype have higher MTHFR activity than 1298AA/677CT (83.2% versus 66.8% relative enzyme function), but also had higher mean homocysteine levels (13.6 µmol/L versus 12.8 µmol/L).^12^ The non-linearity of the association between MTHFR and homocysteine levels is typified by the 1298AA/677TT genotype, who have around 75% loss of enzyme function and 50% higher mean homocysteine levels but, importantly, display a high degree of variability and values that do not appear to be normally distributed. Therefore, any specific recommendations to this group must be based in phenotypic measurements, including individual homocysteine levels and nutrient status. Indeed, though MTHFR is associated with the folate cycle, ensuring adequate B6 and B12 may be at least as important with respect to homocysteine levels.^24^ Homocysteine in 677TT carriers can also be significantly reduced with a small amount of supplemental riboflavin.^25^ This again suggests that phenotypic measurements and ensuring adequate environmental/nutrient status has a much greater impact than does knowledge of genotype. However, it must be cautioned that, as yet, reducing homocysteine with nutritional supplements has not yet been shown to result robustly improve health outcomes, though there may be a small reduction in stroke risk.^26^

This study does have some limitations. The approach used relies on the use of both simulated and statistically-ideal normal distributions based on published descriptive data rather than the data itself. However, where the methods could be tested against known data, such as the degree to which the glucose risk score explains glucose variability, the results were very similar to the original analyses. Importantly, if this approach fails to accurately recreate datasets similar to those in the published literature, then it is likely that those datasets were not normally-distributed and the original analyses were therefore inappropriate. This is probably the case for homocysteine levels in individuals with the MTHFR 1298AA/677TT genotype based on the widely-cited study by van der Put *et al*..^12^ Though all the SNPs analysed here have low penetrance, they were specifically chosen because they are well-characterized in multiple populations and commonly included in third party DTC analyses of consumer genetic data. Though we have only highlighted a few SNPs, the techniques applied here could be used by any practitioner or interested individual to better understand their disease or outcome risk based on common genetic SNPs.

Even though there is inherent error in our approach, it is clear that using population means to determine genetic risk and make recommendations based on genetics, as is very common in the DTC market, is likely to be highly-flawed due to inherent phenotypic variability. This is includes variability in risk based on common factors such as socioeconomic status and ethnicity. For instance, FTO genotypes are associated with increased BMI in Caucasians, but not in those of African origin.^7^ For the risk of both obesity and prediabetes or type 2 diabetes, particularly, the effect of the environment (diet, exercise, nutrient status) is likely to dominate the phenotype such that knowing about an individual’s SNPs associated with risk will have little benefit. A focus on genetic risk may indeed be detrimental due to the fact that i) thinking that you have a risk SNP can have an effect on physiology regardless of whether you have that SNP,^27^ ii) the majority of people have average genetic risk for a given phenotype, iii) DTC genetics testing still includes significant variability and error,^28^ iv) there is little to no evidence that specific interventions for a given common SNP have any effect on health outcomes, v) communicating genetic risk does not appear to alter health behaviours,^29^ and vi) though statistically significant, the final effect of most SNPs on phenotype could often be considered physiologically irrelevant. These risks have generally been acknowledged by the scientific community performing genetic research, but the over-interpretation of risk by third-parties relying on published population averages remains a significant worry, likely due to misinterpretation of the nature of the data.

## CONCLUSIONS

Using simple statistical techniques, either with Python code or freely-available online tools, we have outlined a method by which healthcare providers and third-party genetic analysis tools can more accurately analyse genetic disease risk. Importantly, it is worth noting that the widely-characterized and cited SNPs for obesity, type 2 diabetes, and methylation status appear to have negligible overall effects on phenotype compared to the dominant effect of the environment.

## ACKNOWLEDGEMENTS

T.R.W is supported by start-up funds from the University of Washington Department of Pediatrics.

## AUTHOR CONTRIBUTIONS

T.R.W developed the concept, performed the statistical analyses, and drafted the manuscript. N.O performed the number generation, assisted with the statistical analyses, and edited the manuscript. Both authors approved the final version.

## DATA AVAILIBILITY

All data was randomly-generated based on published descriptive measures from population studies.

